# NGF-mediated photoablation of nociceptors reduces pain behavior in mice

**DOI:** 10.1101/575274

**Authors:** L Nocchi, C Portulano, F Franciosa, B Doleshall, M Panea, N Roy, M Maffei, A Gargano, E Perlas, PA Heppenstall

## Abstract

Nerve growth factor (NGF) and its receptors TrkA and p75 play a key role in the development and function of peripheral nociceptive neurons. Here we describe novel technology to selectively photoablate TrkA positive nociceptors through delivery of a phototoxic agent coupled to an engineered NGF ligand and subsequent near infrared (NIR) illumination. We demonstrate that this approach allows for on demand and localized reversal of pain behaviors in mouse models of acute, inflammatory, neuropathic and joint pain. To target peripheral nociceptors we generated a SNAP-tagged NGF derivative, NGF^R121W^ that binds to TrkA/p75 receptors but does not provoke signaling in TrkA positive cells or elicit pain behaviors in mice. NGF^R121W-SNAP^ was coupled to the photosensitizer IRDye®700DX phthalocyanine (IR700) and injected subcutaneously. Following NIR illumination of the injected area, behavioral responses to nociceptive mechanical and sustained thermal stimuli, but not innocuous stimuli, were substantially reduced. Similarly, in models of inflammatory, osteoarthritic and neuropathic pain, mechanical hypersensitivity was abolished for three weeks following a single treatment regime. We demonstrate that this loss of pain behavior coincides with the retraction of neurons from the skin which then re-innervate the epidermis after 3 weeks corresponding with the return of mechanical hypersensitivity. Thus NGF^R121W-SNAP^-mediated photoablation is a minimally invasive approach to reversibly silence nociceptor input from the periphery, and control pain and hypersensitivity to mechanical stimuli.

## Introduction

Current therapeutic strategies for the management of chronic pain such as nonsteroidal anti-inflammatory drugs (NSAIDs), antidepressants, and opioids [6; 9; 49] have limited efficacy and may be associated with dependence, tolerance, and serious side effects [29; 59]. To avoid these issues, and develop pain therapies which are more efficacious, much effort has been made in developing strategies to target the peripheral nervous system, and in particular molecules which are selectively expressed in nociceptors. Amongst the most promising of these molecules, nerve growth factor (NGF) and its cognate receptors TrkA/p75 have emerged as frontrunners for the design of next generation analgesics [13; 15; 41-43; 53].

There is a wealth of data supporting the role of NGF and its receptors in pain signaling. NGF and TrkA are required for the development of the peripheral nervous system; in their absence, both in animal models, and human genetic disease, nociceptive neurons fail to develop, and animals and human patients display severe deficits in pain sensitivity [11; 21; 27; 28; 32; 50; 56]. In the adult peripheral nervous system, TrkA receptors are expressed predominantly on peptidergic nociceptors, and NGF signaling plays an important role in setting pain sensitivity. Thus cutaneous administration of NGF leads to hyperalgesia within 1–3 hours, both in rodents and humans [20; 33; 34; 47; 52] and endogenous NGF levels are elevated in chronic pain conditions and promote peripheral and central sensitization [36; 40; 48]. The recognition that NGF has a critical role in the generation and potentiation of pain has created a strong rationale for developing methods that interfere with its signaling. The most successful of these approaches has been the development of antibodies against NGF [13] such as tanezumab and fasinumab, which have advanced to phase III trials for osteoarthritis (OA) of knee and hip. There are however a number of safety issues associated with antagonizing NGF, the most prominent of which is a rapidly progressive OA, leading to joint replacement in some trial patients [24].

Rather than inhibiting NGF signaling at the molecular level, here we have asked whether the NGF ligand may offer a means of accessing nociceptive neurons via their TrkA receptors to deliver a phototoxic agent to silence their activity. To test this hypothesis, we generated a self labelling recombinant NGF-SNAP tag fusion protein that could be covalently conjugated to the NIR photosensitizer IRDye®700DX phthalocyanine (IR700). While injection of NGF^SNAP^-IR700 and subsequent NIR illumination reduced nociceptive behavior in mice, results were confounded by the fact that NGF is in itself proalgesic. We thus turned to “painless” mutated derivatives of NGF which still bind to receptors but provoke abrogated signaling, in order to deliver the photosensitizer. Such NGF mutants have been described in patients with Hereditary Sensory and Autonomic Neuropathy type 5 (HSAN V), who display a loss of pain perception and other sensory and autonomic abnormalities [1] as a result of mutations in the *NGF* gene. To date, three mutations in NGF have been identified in HSAN V patients, p.R221W, p.V232fs and p.R121W, that reduce the processing of NGF, alter the neurotrophic activity or abolish the formation of mature NGF-β, respectively [12; 21; 30; 54; 57]. Of these, we selected the NGF p.R121W mutation as a delivery tool as this gave the highest yields in recombinant production when fused to the SNAP tag. We found that NGF^R121W-SNAP^ was able to bind to TrkA positive cells but did not provoke TrkA mediated signaling, or elicit sensitization when injected in mice. Moreover, conjugation of NGF^R121W-SNAP^ to IR700, injection in mice, and subsequent NIR illumination of the injected area led to long term reversal of nociceptive behavior that paralleled retraction of cutaneous nerve fibers from the epidermis. Thus, through ligand guided delivery of a small molecule photosensitizer we are able to achieve selective disruption of nociceptor input, and substantially reduced pain.

## Results

### Generation and characterization of NGF^SNAP^

Recombinant NGF was cloned and purified as a SNAP tag fusion protein (NGF^SNAP^), which allows for covalent coupling to BG-derived photosensitizers and imaging fluorophores. We first tested the binding selectivity of NGF^SNAP^ by labelling it with a BG-fluorophore and applying it to cells expressing neurotrophin receptors. NGF^SNAP^ was coupled in vitro to BG-Surface549, and applied at a 100 nM concentration to Hek293T cells transiently transfected with TrkA, TrkB or TrkC and p75. NGF^SNAP^ bound its cognate receptor, decorating the surface of TrkA/p75 transfected cells (**Fig. 1 a**), while cells transfected with BDNF/NT4 and NT3 receptors with p75 were not stained (**Figs. 1 b, c**), demonstrating that binding occurs faithfully in a receptor-specific fashion.

**Figure 1.**
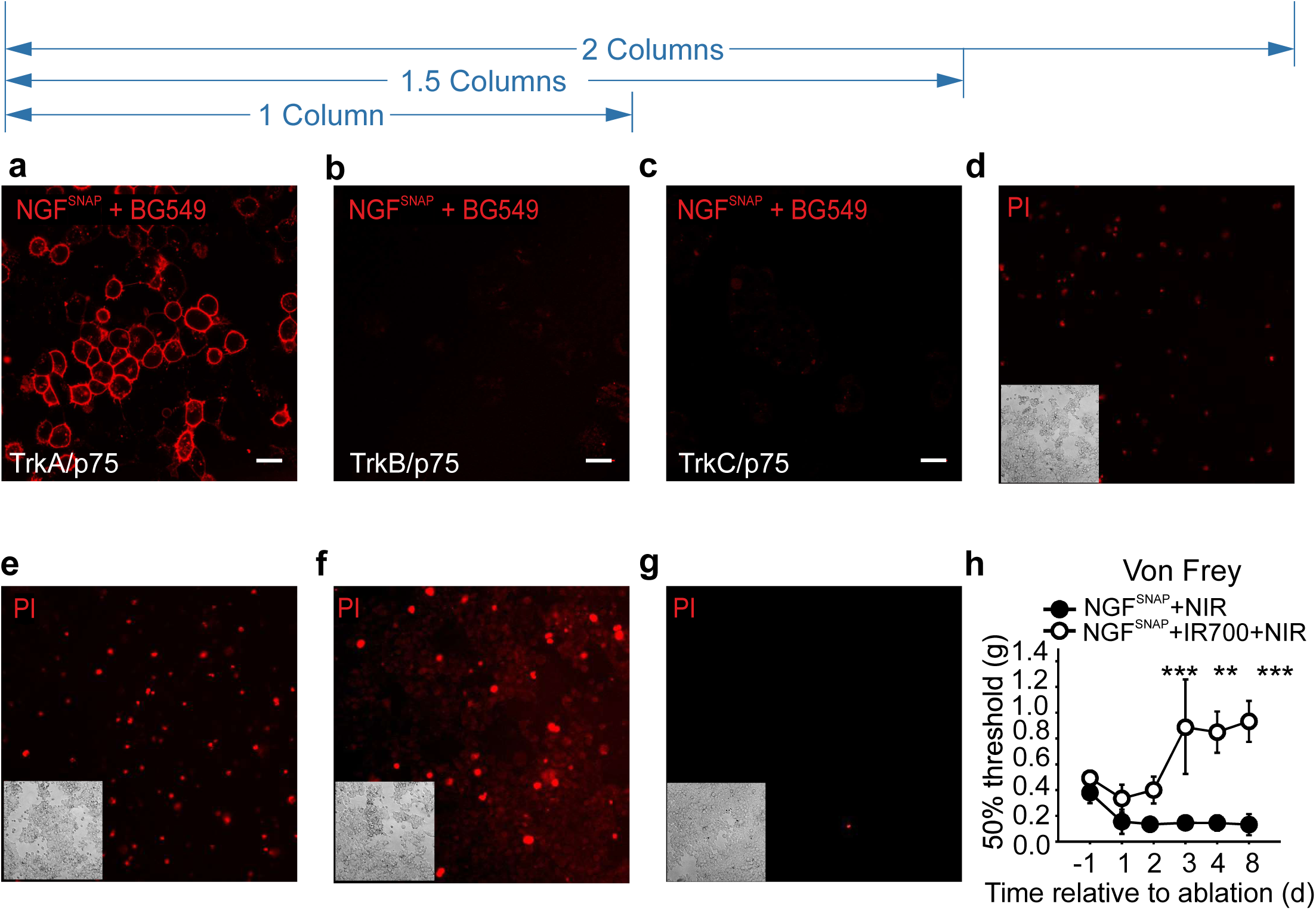
Characterization of NGF^SNAP^. (**a-c**) Confocal images of Hek293T cells transiently transfected with TrkA, TrkB or TrkC (panel **a, b** and **c** respectively) and p75 stained with 100 nM NGF^SNAP^ conjugated to surface BG-546. NGF^SNAP^ selectively binds only to TrkA/p75 transfected cells, with virtually no binding to TrkB or TrkC transfected cells. Scale bar 20 μm. (**d-f**) Hek293T cells transiently transfected with TrkA and p75 were incubated with NGF^SNAP^ (0.1, 0.5 and 1 μM, panel **d**, **e** and **f** respectively) conjugated with BG-IR700, and then exposed to NIR illumination for 2 min. 24 h post *in vitro* photoablation, cells where stained with propidium iodide (PI). 1 μM BG-IR700-conjugated NGF^SNAP^ (panel **g**) was applied onto mock transfected cells, and cells exposed to 690 nm light for 2 min as a negative control. Insets: images of corresponding brightfields. (**h**) Von Frey thresholds at baseline and at each of 3 consecutive days of hind paw injection of NGF^SNAP^ conjugated with BG-IR700 (closed circles) or unconjugated (open circles), followed by NIR light exposure. n=4. Error bars indicate SEM. ***p<0.001; **p=0.01 (two-way ANOVA).

Having confirmed that NGF^SNAP^ specifically labels TrkA expressing cells, we next asked whether a photosensitizer can be coupled to the ligand to ablate TrkA positive cells. Hek293T cells transfected with TrkA/p75 were incubated with NGF^SNAP^ conjugated to the near infrared photosensitizer BG-IR700 and illuminated with NIR light for 2 minutes. 24 hours later cell death was analyzed using propidium iodide staining. In cells transfected with TrkA/p75 we observed concentration dependent increase in cell death upon treatment with NGF^SNAP^ IR700 and NIR illumination (**Figs. 1 d-f**). In the absence of TrkA expression, cell viability was fully preserved following NGF^SNAP^ IR700 application and illumination (**Fig. 1 g**).

We next asked whether NGF^SNAP^ IR700 can be used to photoablate nociceptors in vivo. NGF^SNAP^ conjugated with BG-IR700, (**Fig. 1 h**, filled circles) or unconjugated NGF^SNAP^ (open circles) was injected into the hind paw of C57BL/6J male naïve mice, and skin was illuminated for 2 minutes with IR light exposure on 3 consecutive days. Behavioral responses to calibrated Von Frey filaments were monitored after each ablation at 24 hours and 5 days after the last treatment. As shown in figure 1h, mechanical thresholds increased in ablated mice, suggesting that TrkA expressing nociceptors were targeted by NGF^SNAP^-IR700. However, we also observed a decrease in von Frey thresholds in control NGF^SNAP^ mice (**Fig. 1h**), indicating that the ligand in itself was having a proalgesic effect and might constitute a confounding factor in the measurement of mechanical thresholds in ablated mice. In further experiments, we thus sought to exploit a “painless” NGF mutant described in HSANV patients, NGF^R121W^, which has been described to bind to TrkA receptors but not provoke nociceptive signaling [54].

### Generation and characterization of NGF^R121W-SNAP^

A C-terminal fusion of NGF^R121W^ and SNAP (NGF^R121W-SNAP^) was produced in CHO cells at yields similar to wildtype NGF^SNAP^ and could be readily labelled with BG-549 indicating that the SNAP tag was fully functional **(Fig. 2 a)**. To characterize the binding and signaling properties of NGF^R121W-SNAP^, we first assessed its capacity to label Hek293T cells transfected with neurotrophin receptor plasmids. NGF^R121W-SNAP^ was coupled in vitro with BG-Surface549 and applied to cells. Similar to wildtype NGF^SNAP^, we observed robust membrane labelling of TrkA/p75 expressing cells **(Fig. 2 b)** but no signal in TrkB/p75 or TrkC/p75 transfected cells (**Fig 2 c, d**). We further analyzed downstream signaling provoked by wildtype NGF^SNAP^ and NGF^R121W-SNAP^ by assessing phosphorylation of MAPK and AKT upon treatment of PC12 cells with ligands [22]. Wildtype NGF treatment produced a substantial increase in phosphorylation of MAPK and AKT, while levels of MAPK and AKT phosphorylation in cells treated with NGF^R121W-SNAP^ were no different from untreated cells (**Fig 2 e, f**). We next evaluated the neurotrophic activity of wildtype NGF and NGF^R121W-SNAP^ by quantifying the number of differentiated PC12 cells (assessed by neurite outgrowth [10]) after 6 days of incubation with ligands. A significantly higher proportion of differentiated PC12 cells were present in samples treated with NGF^SNAP^, while those treated with NGF^R121W-^ SNAP were not significantly different from untreated cells (**Fig. 2 g-j**). Finally, we considered the pro-nociceptive activity of NGF^SNAP^ and NGF^R121W-SNAP^ by injecting ligands into the hindpaw of mice and evaluating mechanical and thermal hyperalgesia with the von Frey and hotplate tests. We found that wildtype NGF induced substantial sensitization to both mechanical and thermal stimuli, while injection of NGF^R121W-SNAP^ did not change thresholds to either of these stimuli (**Fig. 2 k, l**). We conclude that, while binding and receptor specificity of NGF^R121W-SNAP^ are comparable to that of NGF^SNAP^, it has the distinctive advantage that it allows for uncoupling of receptor engagement from nociceptive activity, making it a valuable tool to selectively target TrkA expressing peptidergic neurons for the development of an anti-nociceptive treatment strategy.

**Figure 2.**
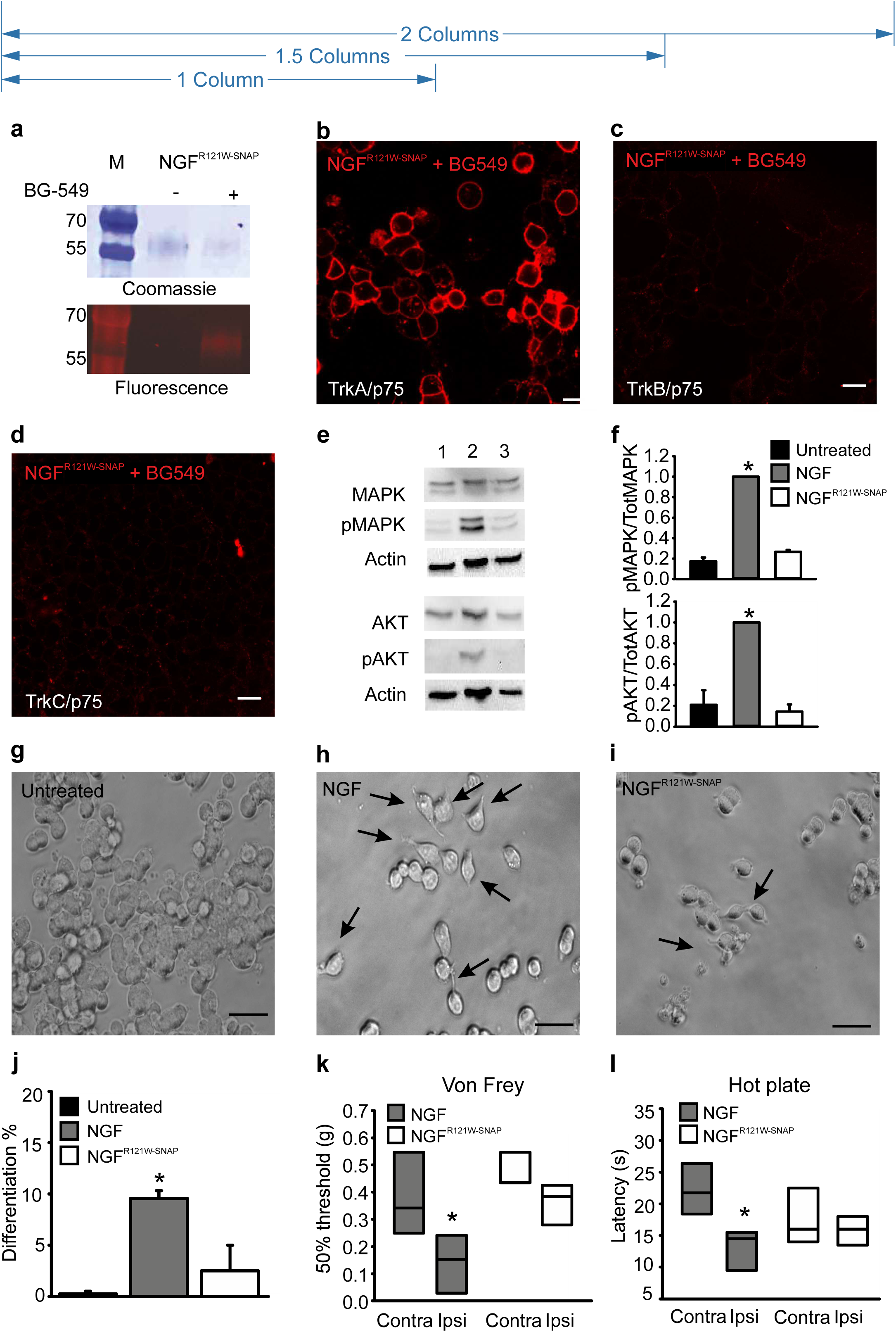
Characterization of NGF^R121W-SNAP^. (**a**) NGF^R121W-SNAP^ was coupled with BG-549 in vitro, run on an SDS-Page, stained with Coomassie blue, and in-gel fluorescence visualized under a UV-light. (**b-d**) Confocal images of Hek293T cells transiently transfected with TrkA, TrkB or TrkC (panel **b, c** and **d** respectively) and p75 stained with 100 nM NGF^R121W-SNAP^ conjugated to surface BG-546. NGF^R121W-SNAP^ selectively binds only to TrkA/p75 transfected cells, with virtually no binding to TrkB or TrkC transfected cells. Scale bar 20 μm. **(e)** Representative western blot of three independent experiments, showing the expression level of MAPK, phospho MAPK, AKT, phospho AKT and actin (loading control) in untreated PC12 cells (lane 1), treated with NGF (lane 2) or NGF^R121W-SNAP^ (lane 3). **(f)** Levels of each protein were expressed as ratio between the phosphorylated form and the total counterpart and then normalized to the NGF-treated sample. (**g-j**) Neuron differentiation in untreated **(g)**, NGF-treated **(h)** and NGF^R121W-SNAP^-treated **(i)** PC12 cells, after 6 days of treatment. Arrows indicate differentiated cells. Scale bar 20 µm. **(j)** Quantitation of neuron-like differentiated PC12 cells, expressed as percentage (%); for the analysis, 241 untreated cells, 246 NGF-treated-cells and 285 NGF^R121W-SNAP^-treated cells were considered. Error bars indicate SEM. **(k)** Von Frey thresholds after injection of NGF (shaded box) or NGF^R121W-SNAP^ (open box) into the hind paw (ipsi) of mice (n=4 animals). Box plots represent the force expressed in grams (g) required to trigger a 50% response. * p=0.05 (two-tailed t-Test). **(l)** Hotplate test after injection of NGF (n=6 animals, red box) and NGF^R121W-SNAP^ (n=5 animals, grey box) into the hind paw of the mice. Box plots represent the latency expressed in seconds (s) of the paw withdrawal in response to heat (52°C). * p=0.009 (two-tailed t-Test).

### NGF^R121W-SNAP^-mediated photoablation and acute nociception

We first assessed the efficacy of NGF^R121W-SNAP^ IR700 mediated photoablation in vitro in Hek293T cells transfected with TrkA/p75. Similar to NGF^SNAP^, the mutant ligand induced a robust and concentration dependent increase in cell death 24 hours after application to cells and NIR illumination (**Fig. 3 a-c**). In contrast, NGF^R121W-SNAP^ IR700 and NIR illumination was without effect in mock transfected cells not expressing TrkA/p75 (**Fig. 3 d**).

**Figure 3.**
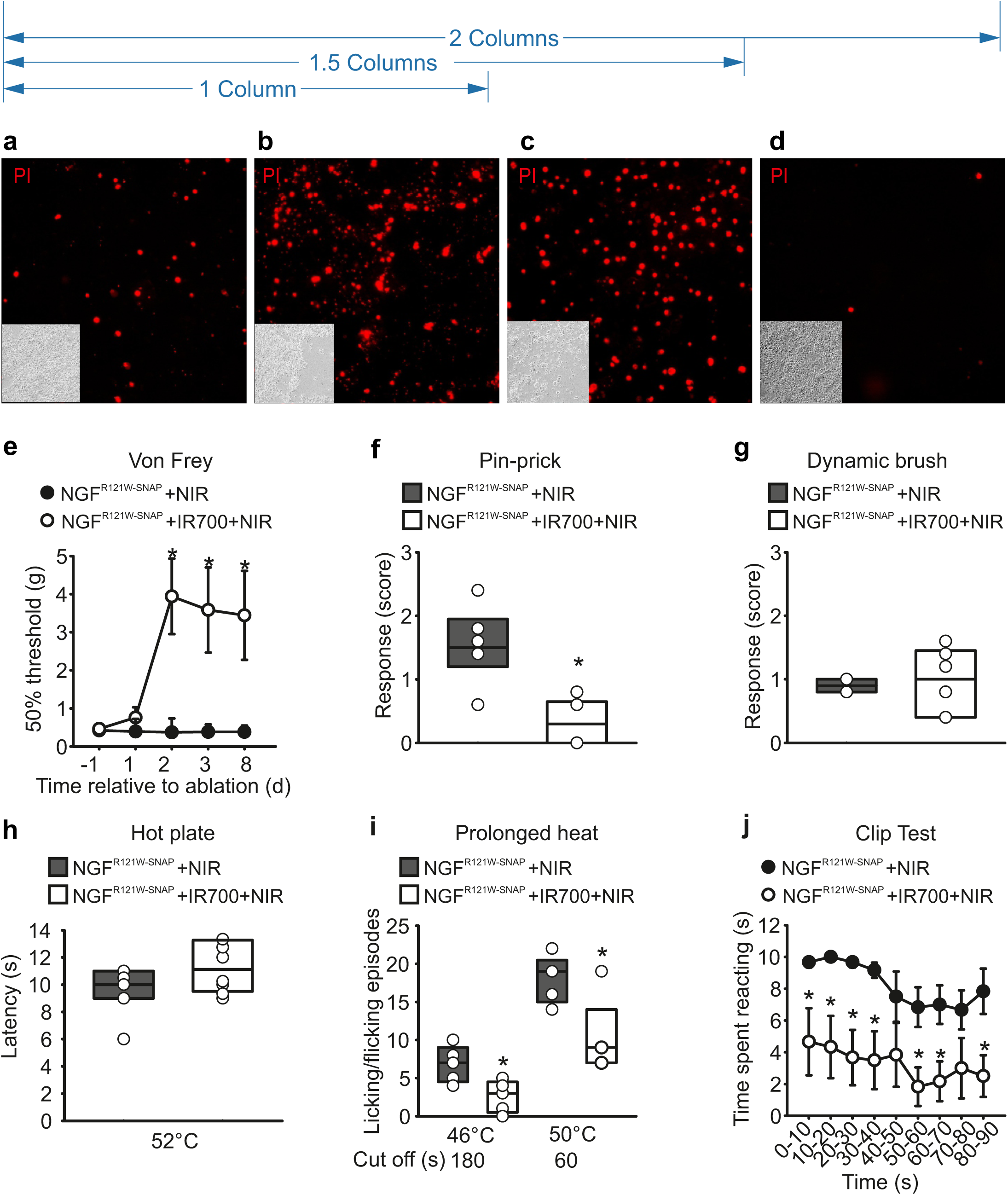
NGF^R121W-SNAP^-mediated ablation in vitro and in acute nociceptive tests. (**a-d**) Hek293T cells transiently transfected with TrkA and p75 were incubated with NGF^R121W-SNAP^ (0.1, 0.5 and 1 μM, panel **a, b** and **c** respectively) conjugated with BG-IR700, and then illuminated with NIR light for 2 min. 24 h post in vitro photoablation, cells where stained with propidium iodide (PI). 1 μM BG-IR700-conjugated NGF^R121W-SNAP^ (panel **d**) was applied onto mock transfected cells, and cells illuminated with 690 nm light for 2 min as a negative control. Insets: images of corresponding brightfields. **(e)** Von Frey thresholds after three days of injection of NGF^R121W-SNAP^ into the hind paw of the mice, with (open circles) or without (closed circles) IR700, followed by NIR illumination (n=6 animals). The test was performed at day 1, 2, 3 and 8 after last ablation treatment. The graph shows the force expressed in grams (g) required to trigger a 50% response. Error bars indicate SEM. *p = 0.001 (two-way ANOVA followed by Bonferroni post hoc test). **(f, g)** Pin-prick and dynamic brush tests after three days of injection of NGF^R121W-SNAP^ into the hind paw of the mice with (open box) or without (shaded box) IR700, followed by NIR illumination (n=6 animals). The test was performed at 7 days after the last ablation treatment. *p = 0.002 (pin-prick, two-tailed t-Test). The graphs show the response to the stimulus, 0-3 score (score=0 no response; score=1 paw withdrawal; score=2 prolonged paw withdrawal; score=3 paw flicking/licking). **(h)** Response latency expressing the time needed to observe a nocifensive behavior upon heat stimulation in a hotplate pre-set at 52°C. Mice were injected for 3 consecutive days with NGF^R121W-SNAP^ into the hind paw, with (open box) or without (shaded box) IR700, followed by NIR illumination (n=6 animals). The test was performed at 7 days after the last ablation treatment. **(i)** Paw licking/flicking behavior evoked by a prolonged heat noxious stimuli using a hotplate (46°C, 50°C) in mice injected in both hind paws for three days with NGF^R121W-SNAP^ with (open boxes) and without (shaded boxes) IR700, followed by NIR illumination (n=5 animals). A cut-off time was set at 180 s (46°C), and 60 s (50°C) *p = 0.018 (two-tailed t-Test). **(j)** Clip tail test after three days of cream application of NGF^R121W-SNAP^ with (n=6 animals, white circles) and without (n=animals, black circles) IR700, followed by NIR illumination on the mouse tail. The test was performed at day 1, 5, 7, 10, 14 and 21 after last ablation treatment. The graph shows the latency expressed in seconds (s) to the clip. Error bars indicate SEM.*p = 0.001 (two-way ANOVA followed by Tukey post hoc test).

To characterize NGF^R121W-SNAP^ mediated photoablation in vivo, we evaluated its efficacy in models of acute nociception. Wild type C57BL/6J male mice were injected for 3 consecutive days in the hind paw with either NGF^R121W-SNAP^, with or without IR700, or with IR700 alone (some controls shown in **Supplementary Fig. 2**) followed by NIR illumination. We first tested mice for sensitivity to punctate mechanical stimuli using von Frey filaments. As shown in **Fig. 3 e**, two days following ablation, mice treated with NGF^R121W-SNAP^ IR700 displayed significantly reduced sensitivity to these stimuli, requiring weights above 3g to provoke a behavioral response, and this insensitivity persisted throughout the 8 day monitoring period. Similarly, responses to nociceptive pin-prick stimuli applied to the paw were also substantially reduced in photoablated compared to control mice (**Fig. 3 f**). We further assessed sensitivity to non-nociceptive dynamic brush stimuli, and observed no effect of NGF^R121W-SNAP^ IR700 (**Fig. 3 g)** indicating that the photoablation is selective for nociceptors. We also tested thermal responsiveness using the hotplate test. Intriguingly, first line reflex latencies were not different between photoablated and control animals, however, licking behavior evoked by sustained noxious thermal stimuli was reduced in treated animals at 46°C and 50°C (**Fig. 3 h, i**). Thus, TrkA positive neurons may mediate pain resulting from tissue damaging thermal stimuli, rather than reflexive defensive responses. Finally, we asked whether NGF^R121W-SNAP^ IR700 can be applied topically to alleviate acute nociception. A micro-emulsion pre-mixed with either NGF^R121W-SNAP^ with or without IR700, or IR700 alone (**Supplementary Fig. 2 e**), was applied to the tail skin and illuminated with NIR light. One week later a calibrated clip was applied to the base of the tail and responses monitored. We observed significantly reduced behavioral responses to the clip in NGF^R121W-SNAP^ IR700 mice throughout the monitoring period, such that mice appeared essentially insensitive to this nociceptive stimulus (**Fig. 3 j**, **Supplementary Video 1**).

### NGF^R121W-SNAP^-mediated photoablation in models of inflammatory, osteoarthritic and neuropathic pain

We investigated the effects of NGF^R121W-SNAP^ mediated photoablation on persistent pain using models of inflammatory, osteoarthritic and neuropathic pain.

To model inflammatory pain, mice were injected in the hind paw with complete Freund’s adjuvant (CFA). After 48 hours, mechanical and thermal hyperalgesia were evident and mice were treated for three days with NGF^R121W-SNAP^ with or without IR700 (**Supplementary Fig. 2 e**), or with NGF^R121W-SNAP^ IR700, with or without near NIR illumination (**Fig. 4 a, b**). Behavioral thresholds to mechanical and thermal stimuli were monitored for 12 days after CFA injection. As shown in **Fig. 4 a**, mice which received NGF^R121W-SNAP^ and NIR inflammation displayed a complete reversal of mechanical hypersensitivity that lasted throughout the 12 day observation period (**Fig. 4 a**). Of note, thermal hyperalgesia induced by CFA was not altered by photoablation (**Fig. 4 b**).

**Figure 4.**
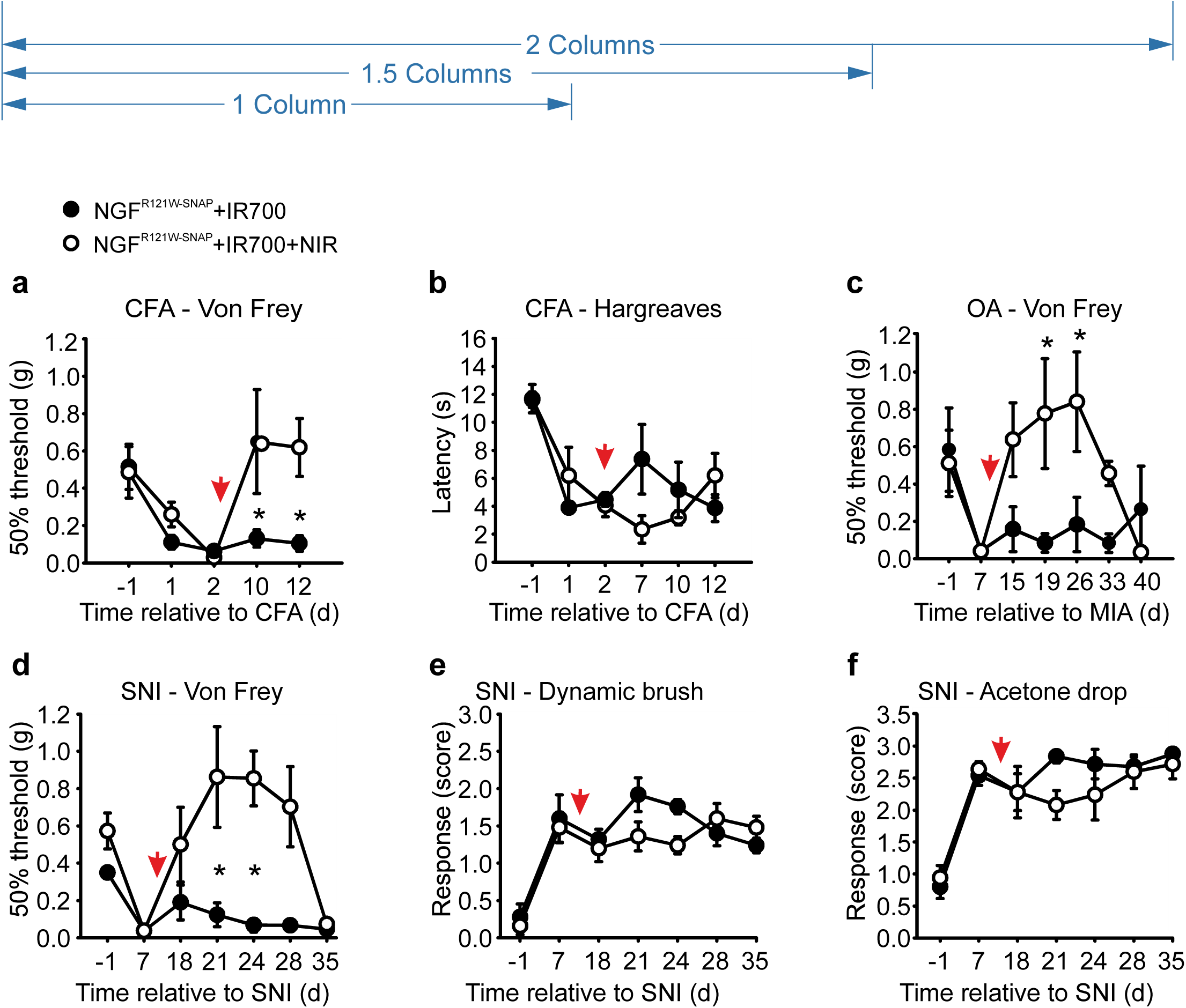
NGF^R121W-SNAP^-guided photoablation in models of prolonged pain. (a, b) Inflammatory pain was measured 24 hour and 48 hours post CFA injection into the paw, and at the indicated days, relative to the CFA injection, after a 3-d course of injections with NGF^R121W-SNAP^ + IR700 with (n=6, open circles) and without (n=6, closed circles) NIR illumination (red arrow indicates the first day of ablation). Baseline before CFA injection is set as d= −1. Error bars indicate SEM. *p = 0.018 (two-way ANOVA followed by Holm Sidak post hoc test). **(c)** Von Frey test in osteoarthritis (OA) mouse model induced by a single injection of monoiodoacetate (MIA) into the knee. The test was performed at the indicated days, relative to MIA injection, and after a 3-d course of injections of NGF^R121W-SNAP^ + IR700 with (n=6, open circles) and without (n=6, closed circles) NIR illumination (red arrow indicates the first day of ablation). Baseline before MIA injection is set as d= −1. Error bars indicate SEM. *p = 0.003 (two-way ANOVA followed by Bonferroni post hoc test). **(d, e, f)** Von Frey, dynamic brush and acetone drop tests in spared nerve injury (SNI) mouse model. The tests were performed at the indicated day relative to the SNI procedure, and after a 3-d course of injections with NGF^R121W-SNAP^ + IR700 with (n=6, open circles) and without (n=6, closed circles) NIR illumination (red arrow indicates the first day of ablation). Baseline before SNI is set as d= −1. Score=0 no response; score=1 paw withdrawal; score=2 prolonged paw withdrawal; score=3 paw flicking/licking; score=4 prolonged paw licking. *p = 0.005 (two-way ANOVA followed by Bonferroni post hoc test).

To model OA-like pain, mice received an intra-articular injection into the knee of monosodium iodoacetate (MIA) [45]. Seven days post injection, significant fibrosis was evident in the joint (**Supplementary Fig. 1 a-f**), and mice developed referred mechanical hypersensitivity to von Frey filaments applied to the plantar surface of the paw (**Supplementary Fig. 1 g)**. NGF^R121W-SNAP^ IR700 was injected into the knee and the area was illuminated with NIR light through the skin for 3 consecutive days. Control mice received NGF^R121W-SNAP^ IR700 but no illumination, or IR700 alone with illumination (**Supplementary Fig. 2 f**). In treated mice, we observed a complete recovery of mechanical thresholds to pre-injury levels, which persisted for 3 weeks before returning to the pre-ablation state (**Fig. 4 c**). Control animals displayed no recovery and remained mechanically hypersensitive throughout the 40 day monitoring period (**Fig. 4 c, Supplementary Fig. 2 f**).

For neuropathic pain, we used the spared nerve injury (SNI) model [17]. 7 days following nerve injury when all mice had developed mechanical hypersensitivity, NGF^R121W-SNAP^ IR700 was injected into the paw and the skin was illuminated with NIR light on 3 consecutive days. Control animals received NGF^R121W-SNAP^ IR700 without illumination (**Fig. 4 d-f**), IR700 with illumination, or NIR illumination alone (**Supplementary Fig. 2 g**). Mice were then assayed for hypersensitivity to mechanical punctate stimuli, dynamic brush stimuli, and evaporative cooling. In mice treated with NGF^R121W-SNAP^ IR700 and NIR illumination we again observed that thresholds to punctate mechanical stimuli recovered to pre-injury levels for about 3 weeks, before returning to a hypersensitive state (**Fig. 4 d**). Intriguingly, responses to dynamic brushing mechanical stimuli were not altered by photoablation (**Fig. 4 e**), nor were responses to evaporative cooling (**Fig. 4 f**).

### Mechanism of NGF^R121W-SNAP^-mediated photoablation

To understand how NGF^R121W-SNAP^ mediated photoablation underlies the loss and then recovery of mechanical sensitivity to nociceptive stimuli in treated mice, we performed a histological analysis of skin at 7 and 28 days post treatment.

We first investigated the degree of cell death of non-neuronal cells in the skin by performing TUNEL staining to detect apoptotic DNA fragmentation. Skin was harvested at 7 and 28 days post treatment in animals injected with NGF^R121W-SNAP^ IR700 and illuminated with NIR light, or at 7 days post treatment in control animals receiving only IR700 and illumination. Sections were then processed for TUNEL staining and immunohistochemistry with an anti-K14 antibody to mark keratinocytes. As shown in **Figures 5 a-c**, very few apoptotic cells could be detected in the skin in all conditions and there was no difference in treated versus control samples (**Fig. 5 a-d**). The number of keratinocytes was also constant across conditions and time points. Thus, at the concentrations and light intensities used here, there is no detectable effect of photoablation on non-neuronal cells in the skin.

**Figure 5.**
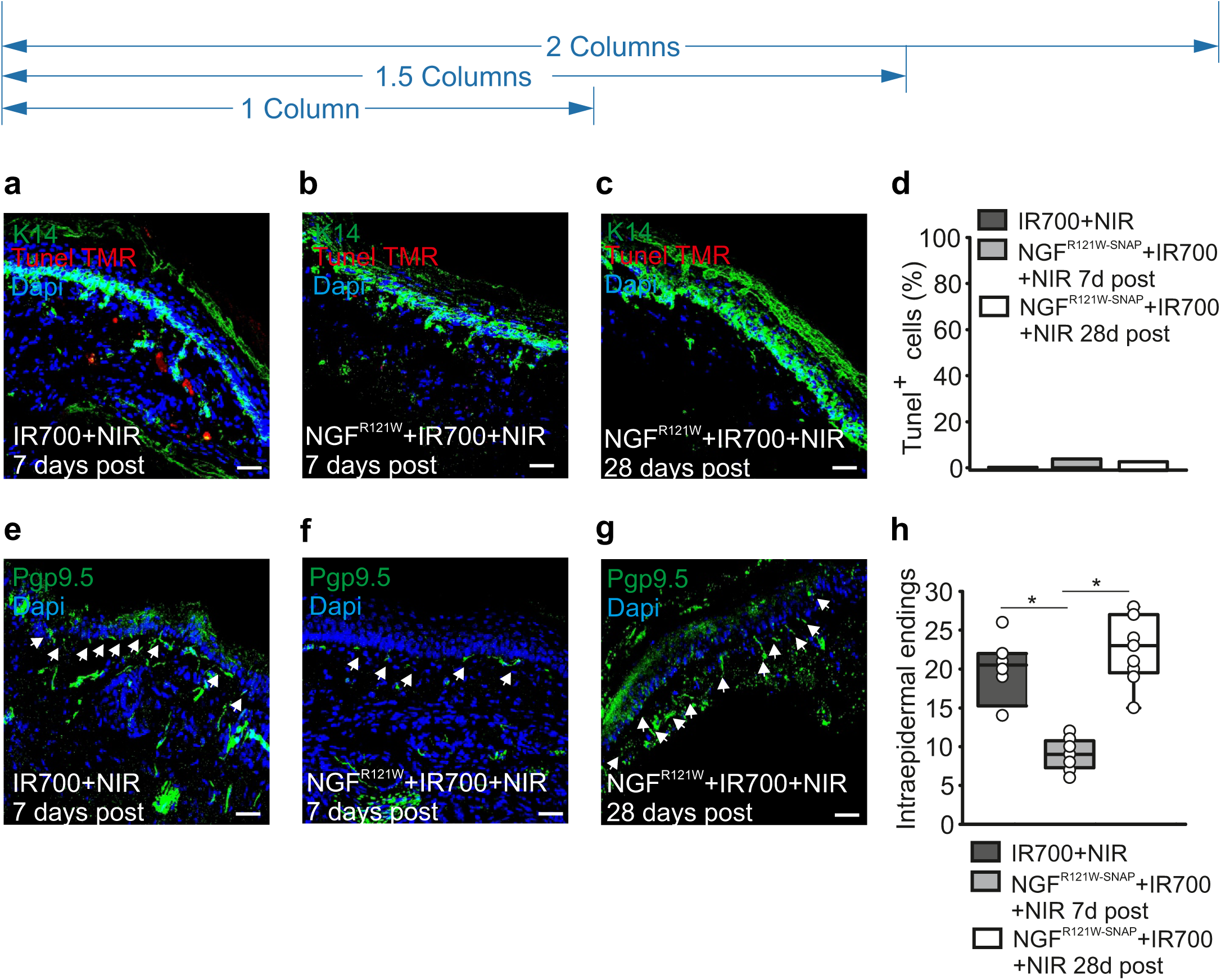
NGF^R121W-SNAP^-guided photoablation induces reversible nerve retraction without affecting keratinocytes in the skin. **(a-d)** Representative paw skin cryosections stained for keratin 14 (K14, in green) and TUNEL-TMR (in red) with quantitation of keratinocytes cell death in mice (n=3 animals) injected for three days with IR700 followed by NIR illumination **(a)**, NGF^R121W-SNAP^+IR700 followed by NIR illumination **(b, c)**. Skin was collected 7 days after **(a, b)** and 28 days after **(c)** the last day of injection. The number of TUNEL^+^ keratinocytes is expressed as percentage **(d)**. **(e-h)** Representative paw skin cryosections stained for the neuronal marker Pgp9.5 (in green) of mice injected for three days with IR700 followed by NIR illumination **(e)**, NGF^R121W-SNAP^+IR700 followed by NIR illumination **(f, g)**. The skin was collected 7 days after **(e, f)** and 28 days after **(g)** the last day of injection. Arrows indicate free nerve endings. Quantitation of the epidermal free nerve endings for the indicated sample; 8 sections were analyzed **(h)**. *p = 0.001 (two-tailed t-Test).

To investigate the effects of NGF^R121W-SNAP^ mediated photoablation on innervation of the skin we performed immunohistochemistry using an anti-Pgp9.5 antibody to detect neuronal fibers. Again, skin samples were taken at 7 and 28 days post treatment in treated animals (NGF^R121W-^ SNAP IR700 plus illumination), or at 7 days post treatment in control animals (IR700 and illumination). We observed a significant reduction in the number of the intra-epidermal free nerve endings in treated animals compared to control at 7 days after injection (**Figs. 5 e, f, h**). Importantly, at 28 days post ablation, the number of fibers was restored to a value similar to the one in the control group (**Figs. 5 g, h**). Thus following photoablation, we observe a retraction of nociceptors from the skin that then reinnervate the epidermis with a similar time course to the loss and return of mechanical sensitivity seen in behavioral experiments.

## Methods

### Animals

Wild type C57BL/6J and Balb/c mice were used at age of 8-10 weeks for all the experiments. All mice were bred and maintained at the EMBL Neurobiology and Epigenetic Unit, Rome, in accordance with Italian legislation (Art. 9, 27. Jan 1992, no 116). Experiments were performed under license from the Italian Ministry of Health, and in compliance with the ARRIVE guidelines.

### Production of recombinant NGF^SNAP^ and NGF^R121W-SNAP^

cDNAs encoding for murine NGF^SNAP^ and NGF^R121W-SNAP^ including a C-terminal poly-histidine tag (His6) inserted for purification purposes were cloned into the pMB-PB vector as a fusion protein [37]. Protein expression was carried out using Chinese hamster ovary (CHO) cells (ATCC, mycoplasma negative) as described by Balasubramanian [2]. Secreted NGF^SNAP^ and NGF^R121W-SNAP^ were purified from cell medium using a Ni-NTA resin (Qiagen, #30210) and eluted with an excess of imidazole. Eluted fractions were then pooled, concentrated, and stored for further analysis.

### Synthesis of BG-IR700

3mg of IRDye®700DX N-hydroxysuccinimide ester fluorophore (LI-COR Biosciences GmbH, Bad Homburg, Germany) were dissolved in 150µl DMSO and treated with 1.5mg BG-PEG11-NH2 and 5µl diisopropylethylamine. After 1h, the product BG-PEG11-IRDye®700DX was purified by HPLC using a Waters Sunfire Prep C18 OBD 5µM; 19 × 150 mm column using 0.1M triethylammonium acetate (TEAA) (pH 7.0) and 0.1M TEAA in water/acetonitrile 3:7 (pH 7.0) as mobile phases A and B, respectively. A linear gradient from 100% A to 100% B within 30 minutes was used. The fractions containing the product were lyophilized. Stock solutions of BG-IR700 were stored at high concentration in DMSO. Prior to conjugation with proteins, BG-IR700 was diluted in physiological buffer.

### *In-vitro* experiments

Hek293T cells were transfected with TrkA, TrkB, TrkC and p75 plasmid using Lipofectamine 2000 (Thermo). For the labelling, 1 µM NGF^R121W-SNAP^ was coupled with 3 µM BG549 surface (NEB # S9112) for 1 hour at 37°C in calcium imaging (CIB) buffer (NaCl 140 mM; KCl 4 mM; CaCl_2_ 2 mM; MgCl_2_ 1 mM; NaOH; 4.55 mM; Glucose 5 mM; HEPES 10 mM; pH 7.4). The coupling reaction was filtered through a PD MiniTrap G-25 column (GE Healthcare #28-9180-07) to remove the excess of BG. Cells were incubated with the coupling reaction for 15-20 minutes at 37°C, then washed 3 times in CIB.

To assess NGF signaling, PC12 cells were cultured on collagen IV-coated 6-well plate in DMEM / F12 medium containing 1% horse serum. Cells were treated for 2 minutes with PBS or 1 µM NGF or 1 µM NGF^R121W-SNAP^, followed by lysis in Ripa buffer.

For the neuronal differentiation assay, PC12 cells were daily exposed to 100 ng/ml NGF (Alomone labs, #N-100) or NGF^R121W-SNAP^ for 6 days. The number of neuron-like cells were counted at day 6.

All images were visualized with a Leica SP5 confocal microscope.

### SDS-Page and Western blot

To assess the coupling reaction, 1 µM NGF^R121W-SNAP^ was coupled with 1.5 µM BG549 for 1 hour at 37°C. The coupling reactions were analyzed by SDS-PAGE on a precast acrylamide gel (BioRad #456-9034), along with the same concentrations of NGF^R121W-SNAP^ alone. The bands corresponding to the binding of NGF^R121W-SNAP^ with BG549 were visualized by gel fluorescence. All the samples were visualized by Coomassie staining.

For NGF-mediated signaling, PC12 cells were collected and lysed in Ripa Buffer (Sigma, #R0278) with proteases inhibitor cocktail (Roche #11873580001). Protein lysates were quantified by Bradford assay. For all the experiments, 30 µg total lysate were separated on 10% SDS-Page gel and transferred to a nitrocellulose membrane (Protran #10600007). Membranes were incubated with the following antibodies: anti MAPK (Cell Signaling #4695), anti phospho MAPK (Thr202/Tyr204) (Cell Signaling #9106), anti AKT (Cell Signaling #4691), anti phospho AKT (Ser473) (Cell Signaling #9271), anti-Actin (Cell signaling #4970). Bands were visualized using the ECL detection system (Amersham #RPN2106); band density was calculated using ImageJ and the levels of phosphorylated proteins were normalized to the total counterpart.

### Immunofluorescence

Skin from the plantar region of the mice hindpaw was dissected 7 days and 28 days after treatment of SNI mice with IR700, with or without NGF^R121W-SNAP^, followed by near IR illumination. Skin was fixed with 4% PFA overnight at 4°C, cryoprotected in 30% sucrose for 6 hours and embedded in Tissue-Tek O.C.T. compound. Twenty-five micrometer thick sections were cut using a cryostat (Leica, CM3050S), permeabilized with 0,3% Triton-X, blocked in 5% goat serum + 0,3% Triton-X and stained for one of the antibodies listed below, prepared in PBS containing 5% goat serum + 0,3% Triton-X. We used K14 (1:200 dilution; Covance # PRB155P) antibody marking for keratinocytes; TUNEL TMR (In situ cell death detection kit – TMR red, Roche # 012156792910) for detecting apoptotic cells; Pgp9.5 (1:200 dilution; Dako # Z5116) marking for nerve fibers. DAPI (1 µg/ml, Invitrogen # D1306) was used to stain nuclei. Slides were mounted with Prolong gold antifade (Invitrogen # P36930), imaged with a Leica SP5 confocal microscope and analyzed with ImageJ. To quantify keratinocytes apoptosis, TUNEL^+^ K14^+^ double positive cells were counted in the epidermal basal layer expressed as percentage. For Pgp9.5^+^ fibers quantitation, the region of dermis / epidermis junction was analyzed and only the intra-epidermal free nerve endings per section were considered for counting [38].

### Knee histology

After intraarticular injection with saline or MIA, knees were dissected out after 7 and 28 days. The surrounding tissues were trimmed and knees were post-fixed overnight and then placed into a decalcifying solution for 72 hours. Decalcified knees were then washed and embedded in paraffin wax. Eight micrometer thick sections were cut and stained for hematoxylin and eosin and for trichrome staining.

### Behavioral testing

All behavior experiments were performed on adult male mice (8-10 week old). Mice were placed on an elevated platform with a mesh floor and habituated for minimum 30 minute, unless otherwise specified.

For the Von Frey test, the plantar side of the hindpaw was stimulated with calibrated von Frey filaments (North Coast medical, #NC12775-99). The 50% paw withdrawal threshold was calculated using the Up-Down method [14]. Post MIA injection, Von Frey testing was confined to the posterior-heel region of the hind paw; post SNI, von Frey testing was confined to the sural nerve innervating region of the paw.

To measure dynamic allodynia, the plantar hind paw was stimulated by stroking using a paint brush (heel-to-toe direction). The responses were scored as previously described [19] as 0 = no response; 1 = brief paw withdrawal; 2 = prolonged paw withdrawal; 3 = paw flicking or licking. Test was performed 5 times, with 1 minute interval between trials.

To measure thermal nociception, mice were placed on a hotplate (Ugo Basile, #35150) preset at 52°C and video recorded. The latency to response (flicking or licking the hindpaw) was measured. In order to avoid tissue damage, a cut off of 30 seconds was defined.

A hotplate was also used to assess the response to prolonged noxious stimuli. The hotplate was set at 46°C and 50°C and different cut offs were set for each temperature: 3 minutes for 46°C, and 1 minute for 50°C, as previously described [25].

Thermal nociception was also evaluated using Hargreaves test, in which a thermal heat stimulus (IR 35) was focused onto the plantar region of the hind paw. To avoid tissue damage, a cut off of 20 seconds was set. The test was repeated 3 times, with 5 minutes interval between trials.

To assess evaporative cold allodynia, the acetone drop test was performed [16]. Briefly, a single drop of cold acetone was applied onto the plantar side of the hindpaw of mice using a syringe without touching the paw of the mice. The responses were scored according to the following scheme: 0 = no response, 1 = paw withdrawal or a single flick, 2 = repeated flicking of the paw and 3 = licking of the paw. The test was repeated five times with 1 minute interval between trials.

For the pin-prick test, the plantar side of the hindpaw was gently touched using an insect pin attached to a 1 g von Frey filament, without penetrating the skin and the yes/no responses were noted for 10 trials, with 1 minute interval between the trials.

For the clip tail test, an alligator clip covered with a rubber casing to reduce the potential tissue damage, was placed on the tail of the mouse, near the base of the tail; then, each mouse was placed on a plexiglass chamber and video recorded. A response was scored when the mice showed a response to the clip by biting, vocalization, grasping of tail or jumping. A cut off of 90 seconds was set and data are plotted as the time to respond (latency) for each mouse.

### Complete Freund’s Adjuvant model

Complete Freund’s Adjuvant (CFA, 20 µl, 2% in 1:1 saline) (Sigma-Aldrich, F5881) was injected into the hindpaw of mice. Behavioral tests were performed at 24h and 48h post CFA injection to assess inflammation, followed by photoablation experiments.

### Spared Nerve Injury model

Peripheral nerve injury was induced by spared nerve injury (SNI) [46]. Briefly, mice were anesthetized using 2.5% isoflurane. The sciatic nerve near the thigh region was exposed and the peroneal and tibial branches were ligated and cut, leaving the sural nerve intact. Behavioral tests were performed starting from 1 week after injury, followed by photoablation experiments.

### Osteoarthritis model

As previously described [45], 10 µl monosodium iodoacetate (MIA, 1mg) (Sigma-Aldrich, I2512) was injected intra-articular, through the intrapatellar ligament, into the knee. Sterile saline was injected in the control mice. Behavioral testing was performed 1 week after the injection.

### *In-vivo* photoablation

The hindpaw, the knee or the tail of the mice were treated for 3 consecutive days with NGF^R121W^-SNAP (5 µM) coupled to BG-IR700 (15 µM) (20 μl and 10 μl for paw and knee injections, respectively, and 20 μl for microemulsion application). Depending on the experiment, the coupling reaction was delivered by injection, or topically using a microemulsion, as described previously [44]. Twenty minutes after delivery, near infrared light at 690nm was applied to the treated skin for 1 minute, at 120-150 J/cm^2^ (PSU-III-FDA diode laser, CNI Laser, China). Behavioral tests were performed starting from 1 day after the last photoablation.

### Statistical analysis

All statistical data are presented as standard error of the mean (SEM) along with the number of samples analyzed (n). Student’s *t-*test and/or analysis of variance ANOVA followed by the appropriate post hoc test were used. Statistical significance was assumed at p<0.05. Sample sizes were determined from the power of the statistical test performed. No animals were excluded and all experiments were performed blinded with order of testing randomized.

## Discussion

Here we describe an approach based on ligand-targeted, light-activated delivery of a phototoxic agent to ablate nociceptors in the skin and reduce pain behavior in mice. To achieve selective nociceptor ablation *in vivo*, we engineered the “painless” HSAN V NGF^R121W^ mutant ligand to specifically deliver a photoactivatable photosensitizer locally to TrkA expressing nociceptors. Upon illumination with NIR light we show that there is a highly effective reversal in pain behavior in mouse models of inflammatory, neuropathic and osteoarthritic pain.

The ligand-mediated photoablation technology we describe here has been previously validated in mice using engineered TrkB and IL31-receptor ligands to control mechanical pain hypersensitivity and inflammatory itch [18; 44]. Based on extensive evidence supporting a role for TrkA signaling in pain [27; 31; 34-36; 39; 56; 62], we selected NGF as a delivery agent to target cutaneous nociceptors in vivo. Following local illumination of the skin, we observed retraction of fibers innervating the epidermis, corresponding to a long lasting but reversible reduction in nociceptive behavior. When the pain behavior returned 3 weeks after photoablation, we also observed a concomitant re-growth of the cutaneous sensory fibers, suggesting that we were targeting nociceptors, the major transducers of pain. Importantly, no damage to other cellular types was detected under the conditions used, despite the fact that TrkA receptors are expressed on keratinocytes and other cells in the skin.[7; 36; 55] This is in agreement with our previous study using interleukin-31 (IL-31) as a ligand to target pruriceptors in the skin. Like NGF receptors, IL-31 receptors are highly expressed in keratinocytes, however we found that IL-31-guided photoablation induced keratinocyte cell death only at very high concentrations, and at much higher light intensities than those used here [44]. This would suggest that neurons are more sensitive to IR700 induced photoablation than other cells in the skin.

Several recent studies have described naturally occurring mutations in the *NGF* gene in patients affected by a rare form of congenital insensitivity to pain, HSAN V. The variants pro-NGF^R221W^ (corresponding to the mature NGF^R100W^) and NGF^R121W^ have been characterized at the protein level [10; 54; 57], and it has been demonstrated that both retain the binding to their receptors TrkA and p75, while downstream signaling (auto phosphorylation of TrkA and phosphorylation of MAPK) is impaired, preventing the nociceptive activity typically exerted by NGF. The identification of these “painless” NGF forms opens up the possibility that these mutated ligands may be exploited as drug delivery agents that bind to nociceptors but do not activate them. We explored this possibility here by fusing a SNAP-tag to NGF^R121W^, and using it to target the small molecule photosensitizer IR700 to nociceptors. Further development of this approach may also allow for the conjugation of other biologically active small molecules to NGF^R121W^ to deliver them into nociceptors in vivo and directly interfere with neuronal activity.

Previous work has shown that selective antagonism of endogenous NGF is highly effective in animal models of many acute and chronic pain states [23; 39; 51]. Among several agents developed to counteract NGF-mediated sensitization, particular attention has focused on monoclonal antibodies such as tanezumab and fasinumab, which have proven effective in the management of osteoarthritic pain [5; 8]. Although these anti-NGF antibodies showed greater efficacy in chronic pain compared to the nonsteroidal anti-inflammatory drugs (NSAIDs), serious adverse effects, including osteonecrosis, neurogenic arthropathy, and morphologic changes in the sympathetic nervous system were reported [4], leading the US Food and Drug Administration (FDA) to place a hold from 2010 to 2015 on clinical studies involving this class of drugs. The reported adverse effects were similar across anti-NGF monoclonal antibodies, suggesting that they are “class-specific effects” [3]. While new evidence is generated in ongoing phase III clinical trials involving tanezumab and fasinumab in OA, the current evaluation of risk/benefit ratio of anti-NGF therapies remains challenging [5].

Similar safety concerns may also be pertinent to NGF-mediated photoablation. However, there are a number of important differences in the photoablation approach compared to monoclonal-anti-NGF-based strategies for treating pain. Firstly, photoablation targets cells rather than one specific molecular mediator of pain, enabling the treatment to bypass the enormous molecular complexity that underlies pain, and to circumvent molecular redundancy, a major issue in developing new analgesic drugs. Secondly, the photoablation is local and can be applied on demand. Thus dosage, light intensity, and frequency of application may constitute useful checkpoints for patient-personalized modulation. Compared to approaches based on monoclonal antibodies which are administered systemically, this may reduce off-target effects on systems other than nociception. Thirdly, our data indicate that a single treatment regime of photoablation is sufficient to obtain analgesia that lasts for several weeks, rather than hours or days, potentially reducing toxicity. Indeed, it may also be possible to increase the effect duration further by performing photoablation in the nerve innervating the injured area, or even to selectively ablate nociceptors permanently in the DRG. Future work, will explore these possibilities and also establish the safety and efficacy of repeated cycles of photoablation.

We found that photoablation using NGF^R121W-SNAP^ was most effective at reducing responses to punctate nociceptive mechanical stimuli. Thus we observed substantially increased von Frey thresholds under basal, inflammatory, osteoarthritic and neuropathic states, and decreased sensitivity to painful clip and pinprick. Of note, behavioral responses to dynamic mechanical stimuli were not altered by photoablation. In previous work we have demonstrated that these dynamic stimuli evoke nociceptive reflexes through TrkB positive mechanoreceptors after nerve injury, and that photoablation using BDNF^SNAP^ is an effective means of reducing these responses [18]. As well as demonstrating that NGF^R121W-SNAP^ and BDNF^SNAP^ mediated photoablation is selective, these data highlight the complexity of sensory processing under different conditions, and indicate that solely targeting nociceptors may not be an optimal strategy in certain pain states.

Given the prominent role of NGF in thermal hyperalgesia, we expected to see a reduction in thermal nociceptive withdrawal latencies in photoablation-treated mice. For example, using a 52°C stimulus we observed similar latencies between ablated and non-ablated mice in acute tests. Similarly, photoablation failed to revert the reduction in latencies induced in a mouse model of inflammatory pain. This apparent discrepancy could be explained by the fact that residual heat-sensitive cutaneous sensory fibers, not expressing TrkA are preserved by the photoablation and could mediate thermal nociceptive reflexes. Indeed, in a recent study, it has been shown that acute noxious heat sensing in mice depends on a triad of transient receptor potential ion channels: TRPM3, TRPV1, and TRPA1 [60], and importantly, none of these channels are expressed exclusively in TrkA positive neurons [61], [58] [26]. Further work using ligands to target other populations of nociceptors for photoablation may uncover the role of these neurons in mediating thermal nociceptive withdrawal reflexes.

While we saw no difference in withdrawal reflexes to heat, we did observe a significant reduction in nocifensive behavior upon prolonged thermal stimulation at 46°C and 50°C. It has recently been demonstrated that these two behaviors are driven by distinct inputs from the periphery and the spinal cord to the thalamus [25]. Indeed, in the periphery, it was shown that non-peptidergic MRGPRD positive neurons underlie reflexive responses that limit injury, while TRPV1 positive nociceptors play a role in behavioral responses to prolonged stimuli and the coping response to pain [25]. It is thus tempting to speculate that NGF^R121W-SNAP^ mediated photoablation selectively targets those neurons that input the “suffering” quality of pain, while sparing the first-line defensive neurons. Such qualities would be optimal for analgesic therapy, as the protective aspect of the peripheral nervous system to noxious heat is preserved, while the perceived pain of the burn is reduced.

## Acknowledgements

We acknowledge the assistance of David Hacker, Laurence Durrer, and Soraya Quinche of the Protein Production and Structure Core Facility of the EPFL in generation of NGF^SNAP^ and NGF^R121W-SNAP^. We also thank Violetta Paribeni and Valerio Rossi for the technical support provided with the animals. This work was funded by EMBL.

## Competing Interests

The authors declare no competing financial and non-financial interest.

---

**Supplementary Figure 1. Characterization of the osteoarthritis (OA) model.** (**a-c**) Hematoxylin and eosin (H&E) and (**d-f**) trichrome staining on paraffin-embedded knee sections (n=10-12 sections) of three representative animals injected with saline (**a**) or MIA (**b-c**) into the knee and then processed after 7 or 28 days. At 28 days post MIA injection, a reduction in the amount of cells (chondrocytes) within the articular cartilage was assessed as loss of nuclear staining with H&E (indicated with the rectangular area). At the same time point, trichrome staining highlights a dramatic increase in fibrotic cartilage/collagen tissue (blue staining). (**g**) Von Frey test reveals increased mechanical allodynia starting 7 days post MIA injection and lasting throughout the 28 day period of observation.

**Supplementary Figure 2. Photoablation controls.** (**a**) Von Frey thresholds after three days of paw injection of NGF^R121W-SNAP^ + IR700 (closed circle) or IR700 alone (open circles), followed by NIR illumination (n=6 animals). The test was performed at day 1, 2, 3 and 8 after last ablation treatment. The graph shows the force expressed in grams (g) required to trigger a 50% response. Error bars indicate SEM. *p = 0.001 (two-way ANOVA followed by Bonferroni post hoc test). (**b, c**) Pin-prick and dynamic brush tests after three days of paw injection of NGF^R121W-SNAP^ + IR700 (open box) or IR700 alone (shaded box), followed by NIR illumination (n=6 animals). The test was performed 7 days after the last ablation treatment. *p = 0.002 (pin-prick, two-tailed t-Test). The graphs show the response to the stimulus, 0-3 score (score=0 no response; score=1 paw withdrawal; score=2 prolonged paw withdrawal; score=3 paw flicking/licking). (**d**) Clip tail test after three days of cream application of NGF^R121W-SNAP^ + IR700 (closed circles) or IR700 alone (open circles), followed by NIR illumination (n=6 animals). The test was performed at day 1, 5, 7, 10, 14 and 21 after the last ablation treatment. The graph shows the latency expressed in seconds (s) to reaction to the clip. Error bars indicate SEM. *p < 0.001 (two-way ANOVA followed by Tukey post hoc test). (**e**) Von Frey test after 24 hour and 48 hours post CFA injection into the paw, to assess inflammatory pain. The test was performed at the indicated day, relative to the CFA injection, and after a 3-d course of injections with NGF^R121W-SNAP^ + IR700 (n=6, closed circles) or IR700 alone (n=6, open circles) followed by NIR illumination (red arrow indicates the first day of ablation). Baseline before CFA injection is set as d= −1. Error bars indicate SEM. *p < 0.001 (two-way ANOVA followed by Holm Sidak post hoc test). (**f**) Von Frey test in OA mouse model induced by a single injection of monoiodoacetate (MIA) into the knee. The test was performed at the indicated day relative to MIA injection and after a 3-d course of injection of NGF^R121W-SNAP^ + IR700 (n=6, open circles) or IR700 alone (n=6, closed circles) followed by NIR illumination (red arrow indicates the first day of treatment). Baseline before MIA injection is set as d= −1. Error bars indicate SEM. *p < 0.001 (two-way ANOVA followed by Bonferroni post hoc test). (**g**) Von Frey test in spared nerve injury (SNI) mouse model. The test was performed at the indicated day relative to the SNI procedure and after a 3-d course of NIR illumination (red arrow indicates the first day of treatment) alone (n=6, open circles) or with injection of IR700 (n=6, closed circles). Baseline before SNI is set as d=-1.

## References

[1] Axelrod FB, Hilz MJ. Inherited autonomic neuropathies. Semin Neurol 2003;23(4):381–390.

[2] Balasubramanian S, Matasci M, Kadlecova Z, Baldi L, Hacker DL, Wurm FM. Rapid recombinant protein production from piggyBac transposon-mediated stable CHO cell pools. J Biotechnol 2015;200:61–69.

[3] Bannwarth B, Kostine M. Targeting nerve growth factor (NGF) for pain management: what does the future hold for NGF antagonists? Drugs 2014;74(6):619–626.

[4] Belanger P, Butler P, Butt M, Bhatt S, Foote S, Shelton D, Evans M, Arends R, Hurst S, Okerberg C, Cummings T, Potter D, Steidl-Nichols J, Zorbas M. From the Cover: Evaluation of the Effects of Tanezumab, a Monoclonal Antibody Against Nerve Growth Factor, on the Sympathetic Nervous System in Adult Cynomolgus Monkeys (Macaca fascicularis): A Stereologic, Histomorphologic, and Cardiofunctional Assessment. Toxicol Sci 2017;158(2):319–333.

[5] Berenbaum F. Targeting nerve growth factor to relieve pain from osteoarthritis: What can we expect? Joint Bone Spine 2018.

[6] Bhangare KP, Kaye AD, Knezevic NN, Candido KD, Urman RD. An Analysis of New Approaches and Drug Formulations for Treatment of Chronic Low Back Pain. Anesthesiol Clin 2017;35(2):341–350.

[7] Botchkarev VA, Yaar M, Peters EM, Raychaudhuri SP, Botchkareva NV, Marconi A, Raychaudhuri SK, Paus R, Pincelli C. Neurotrophins in skin biology and pathology. J Invest Dermatol 2006;126(8):1719–1727.

[8] Bramson C, Herrmann DN, Carey W, Keller D, Brown MT, West CR, Verburg KM, Dyck PJ. Exploring the role of tanezumab as a novel treatment for the relief of neuropathic pain. Pain Med 2015;16(6):1163–1176.

[9] Busse JW, Wang L, Kamaleldin M, Craigie S, Riva JJ, Montoya L, Mulla SM, Lopes LC, Vogel N, Chen E, Kirmayr K, De Oliveira K, Olivieri L, Kaushal A, Chaparro LE, Oyberman I, Agarwal A, Couban R, Tsoi L, Lam T, Vandvik PO, Hsu S, Bala MM, Schandelmaier S, Scheidecker A, Ebrahim S, Ashoorion V, Rehman Y, Hong PJ, Ross S, Johnston BC, Kunz R, Sun X, Buckley N, Sessler DI, Guyatt GH. Opioids for Chronic Noncancer Pain: A Systematic Review and Meta-analysis. JAMA 2018;320(23):2448–2460.

[10] Capsoni S, Covaceuszach S, Marinelli S, Ceci M, Bernardo A, Minghetti L, Ugolini G, Pavone F, Cattaneo A. Taking pain out of NGF: a “painless” NGF mutant, linked to hereditary sensory autonomic neuropathy type V, with full neurotrophic activity. PLoS One 2011;6(2):e17321.

[11] Carroll SL, Silos-Santiago I, Frese SE, Ruit KG, Milbrandt J, Snider WD. Dorsal root ganglion neurons expressing trk are selectively sensitive to NGF deprivation in utero. Neuron 1992;9(4):779–788.

[12] Carvalho OP, Thornton GK, Hertecant J, Houlden H, Nicholas AK, Cox JJ, Rielly M, Al-Gazali L, Woods CG. A novel NGF mutation clarifies the molecular mechanism and extends the phenotypic spectrum of the HSAN5 neuropathy. J Med Genet 2011;48(2):131–135.

[13] Chang DS, Hsu E, Hottinger DG, Cohen SP. Anti-nerve growth factor in pain management: current evidence. J Pain Res 2016;9:373–383.

[14] Chaplan SR, Bach FW, Pogrel JW, Chung JM, Yaksh TL. Quantitative assessment of tactile allodynia in the rat paw. J Neurosci Methods 1994;53(1):55–63.

[15] Chi KR. Outlook for NGF inhibitor painkiller class brightens. Nat Biotechnol 2016;34(7):679–680.

[16] Choi Y, Yoon YW, Na HS, Kim SH, Chung JM. Behavioral signs of ongoing pain and cold allodynia in a rat model of neuropathic pain. Pain 1994;59(3):369–376.

[17] Decosterd I, Woolf CJ. Spared nerve injury: an animal model of persistent peripheral neuropathic pain. Pain 2000;87(2):149–158.

[18] Dhandapani R, Arokiaraj CM, Taberner FJ, Pacifico P, Raja S, Nocchi L, Portulano C, Franciosa F, Maffei M, Hussain AF, de Castro Reis F, Reymond L, Perlas E, Garcovich S, Barth S, Johnsson K, Lechner SG, Heppenstall PA. Control of mechanical pain hypersensitivity in mice through ligand-targeted photoablation of TrkB-positive sensory neurons. Nat Commun 2018;9(1):1640.

[19] Duan B, Cheng L, Bourane S, Britz O, Padilla C, Garcia-Campmany L, Krashes M, Knowlton W, Velasquez T, Ren X, Ross S, Lowell BB, Wang Y, Goulding M, Ma Q. Identification of spinal circuits transmitting and gating mechanical pain. Cell 2014;159(6):1417–1432.

[20] Dyck PJ, Peroutka S, Rask C, Burton E, Baker MK, Lehman KA, Gillen DA, Hokanson JL, O’Brien PC. Intradermal recombinant human nerve growth factor induces pressure allodynia and lowered heat-pain threshold in humans. Neurology 1997;48(2):501–505.

[21] Einarsdottir E, Carlsson A, Minde J, Toolanen G, Svensson O, Solders G, Holmgren G, Holmberg D, Holmberg M. A mutation in the nerve growth factor beta gene (NGFB) causes loss of pain perception. Hum Mol Genet 2004;13(8):799–805.

[22] Greene LA, Tischler AS. Establishment of a noradrenergic clonal line of rat adrenal pheochromocytoma cells which respond to nerve growth factor. Proc Natl Acad Sci U S A 1976;73(7):2424–2428.

[23] Hefti FF, Rosenthal A, Walicke PA, Wyatt S, Vergara G, Shelton DL, Davies AM. Novel class of pain drugs based on antagonism of NGF. Trends Pharmacol Sci 2006;27(2):85–91.

[24] Hochberg MC. Serious joint-related adverse events in randomized controlled trials of anti-nerve growth factor monoclonal antibodies. Osteoarthritis Cartilage 2015;23 Suppl 1:S18–21.

[25] Huang T, Lin SH, Malewicz NM, Zhang Y, Zhang Y, Goulding M, LaMotte RH, Ma Q. Identifying the pathways required for coping behaviours associated with sustained pain. Nature 2019;565(7737):86–90.

[26] Ikeda-Miyagawa Y, Kobayashi K, Yamanaka H, Okubo M, Wang S, Dai Y, Yagi H, Hirose M, Noguchi K. Peripherally increased artemin is a key regulator of TRPA1/V1 expression in primary afferent neurons. Mol Pain 2015;11:8.

[27] Indo Y, Tsuruta M, Hayashida Y, Karim MA, Ohta K, Kawano T, Mitsubuchi H, Tonoki H, Awaya Y, Matsuda I. Mutations in the TRKA/NGF receptor gene in patients with congenital insensitivity to pain with anhidrosis. Nat Genet 1996;13(4):485–488.

[28] Johnson EM, Jr., Gorin PD, Brandeis LD, Pearson J. Dorsal root ganglion neurons are destroyed by exposure in utero to maternal antibody to nerve growth factor. Science 1980;210(4472):916–918.

[29] Kalso E, Aldington DJ, Moore RA. Drugs for neuropathic pain. BMJ 2013;347:f7339.

[30] Larsson E, Kuma R, Norberg A, Minde J, Holmberg M. Nerve growth factor R221W responsible for insensitivity to pain is defectively processed and accumulates as proNGF. Neurobiol Dis 2009;33(2):221–228.

[31] Lewin GR, Lechner SG, Smith ES. Nerve growth factor and nociception: from experimental embryology to new analgesic therapy. Handb Exp Pharmacol 2014;220:251–282.

[32] Lewin GR, Mendell LM. Nerve growth factor and nociception. Trends Neurosci 1993;16(9):353–359.

[33] Lewin GR, Ritter AM, Mendell LM. Nerve growth factor-induced hyperalgesia in the neonatal and adult rat. J Neurosci 1993;13(5):2136–2148.

[34] Lewin GR, Rueff A, Mendell LM. Peripheral and central mechanisms of NGF-induced hyperalgesia. Eur J Neurosci 1994;6(12):1903–1912.

[35] Lindsay RM, Harmar AJ. Nerve growth factor regulates expression of neuropeptide genes in adult sensory neurons. Nature 1989;337(6205):362–364.

[36] Mantyh PW, Koltzenburg M, Mendell LM, Tive L, Shelton DL. Antagonism of nerve growth factor-TrkA signaling and the relief of pain. Anesthesiology 2011;115(1):189–204.

[37] Matasci M, Baldi L, Hacker DL, Wurm FM. The PiggyBac transposon enhances the frequency of CHO stable cell line generation and yields recombinant lines with superior productivity and stability. Biotechnol Bioeng 2011;108(9):2141–2150.

[38] McArthur JC, Stocks EA, Hauer P, Cornblath DR, Griffin JW. Epidermal nerve fiber density: normative reference range and diagnostic efficiency. Arch Neurol 1998;55(12):1513–1520.

[39] McMahon SB. NGF as a mediator of inflammatory pain. Philos Trans R Soc Lond B Biol Sci 1996;351(1338):431–440.

[40] Mendell LM, Albers KM, Davis BM. Neurotrophins, nociceptors, and pain. Microsc Res Tech 1999;45(4-5):252–261.

[41] Miller RE, Block JA, Malfait AM. Nerve growth factor blockade for the management of osteoarthritis pain: what can we learn from clinical trials and preclinical models? Curr Opin Rheumatol 2017;29(1):110–118.

[42] Mullard A. Drug developers reboot anti-NGF pain programmes. Nat Rev Drug Discov 2015;14(5):297–298.

[43] Mullard A. Painkilling anti-NGF antibodies stage phase III comeback. Nat Rev Drug Discov 2018;17(10):697.

[44] Nocchi L, Roy N, D’Attilia M, Dhandapani R, Maffei M, Traista A, Castaldi L, Perlas E, Chadick CH, Heppenstall PA. Interleukin-31-mediated photoablation of pruritogenic epidermal neurons reduces itch-associated behaviours in mice. Nature Biomedical Engineering 2019;3(2):114–125.

[45] Ogbonna AC, Clark AK, Malcangio M. Development of monosodium acetate-induced osteoarthritis and inflammatory pain in ageing mice. Age (Dordr) 2015;37(3):9792.

[46] Pertin M, Gosselin RD, Decosterd I. The spared nerve injury model of neuropathic pain. Methods Mol Biol 2012;851:205–212.

[47] Petty BG, Cornblath DR, Adornato BT, Chaudhry V, Flexner C, Wachsman M, Sinicropi D, Burton LE, Peroutka SJ. The effect of systemically administered recombinant human nerve growth factor in healthy human subjects. Ann Neurol 1994;36(2):244–246.

[48] Pezet S, McMahon SB. Neurotrophins: mediators and modulators of pain. Annu Rev Neurosci 2006;29:507–538.

[49] Riediger C, Schuster T, Barlinn K, Maier S, Weitz J, Siepmann T. Adverse Effects of Antidepressants for Chronic Pain: A Systematic Review and Meta-analysis. Front Neurol 2017;8:307.

[50] Ritter AM, Lewin GR, Kremer NE, Mendell LM. Requirement for nerve growth factor in the development of myelinated nociceptors in vivo. Nature 1991;350(6318):500–502.

[51] Ro LS, Chen ST, Tang LM, Jacobs JM. Effect of NGF and anti-NGF on neuropathic pain in rats following chronic constriction injury of the sciatic nerve. Pain 1999;79(2-3):265–274.

[52] Rukwied R, Mayer A, Kluschina O, Obreja O, Schley M, Schmelz M. NGF induces noninflammatory localized and lasting mechanical and thermal hypersensitivity in human skin. Pain 2010;148(3):407–413.

[53] Schnitzer TJ, Marks JA. A systematic review of the efficacy and general safety of antibodies to NGF in the treatment of OA of the hip or knee. Osteoarthritis Cartilage 2015;23 Suppl 1:S8–17.

[54] Shaikh SS, Nahorski MS, Woods CG. A third HSAN5 mutation disrupts the nerve growth factor furin cleavage site. Mol Pain 2018;14:1744806918809223.

[55] Shibayama E, Koizumi H. Cellular localization of the Trk neurotrophin receptor family in human non-neuronal tissues. Am J Pathol 1996;148(6):1807–1818.

[56] Smeyne RJ, Klein R, Schnapp A, Long LK, Bryant S, Lewin A, Lira SA, Barbacid M. Severe sensory and sympathetic neuropathies in mice carrying a disrupted Trk/NGF receptor gene. Nature 1994;368(6468):246–249.

[57] Sung K, Ferrari LF, Yang W, Chung C, Zhao X, Gu Y, Lin S, Zhang K, Cui B, Pearn ML, Maloney MT, Mobley WC, Levine JD, Wu C. Swedish Nerve Growth Factor Mutation (NGF(R100W)) Defines a Role for TrkA and p75(NTR) in Nociception. J Neurosci 2018;38(14):3394–3413.

[58] Usoskin D, Furlan A, Islam S, Abdo H, Lonnerberg P, Lou D, Hjerling-Leffler J, Haeggstrom J, Kharchenko O, Kharchenko PV, Linnarsson S, Ernfors P. Unbiased classification of sensory neuron types by large-scale single-cell RNA sequencing. Nat Neurosci 2015;18(1):145–153.

[59] Vadivelu N, Gowda AM, Urman RD, Jolly S, Kodumudi V, Maria M, Taylor R, Jr., Pergolizzi JV, Jr. Ketorolac tromethamine - routes and clinical implications. Pain Pract 2015;15(2):175–193.

[60] Vandewauw I, De Clercq K, Mulier M, Held K, Pinto S, Van Ranst N, Segal A, Voet T, Vennekens R, Zimmermann K, Vriens J, Voets T. A TRP channel trio mediates acute noxious heat sensing. Nature 2018;555(7698):662–666.

[61] Vriens J, Owsianik G, Hofmann T, Philipp SE, Stab J, Chen X, Benoit M, Xue F, Janssens A, Kerselaers S, Oberwinkler J, Vennekens R, Gudermann T, Nilius B, Voets T. TRPM3 is a nociceptor channel involved in the detection of noxious heat. Neuron 2011;70(3):482–494.

[62] Woolf CJ, Safieh-Garabedian B, Ma QP, Crilly P, Winter J. Nerve growth factor contributes to the generation of inflammatory sensory hypersensitivity. Neuroscience 1994;62(2):327–331.

