## Supplementary figures and images for "NGF-mediated photoablation of nociceptors reduces pain behavior in mice"

### Suppl fig 1

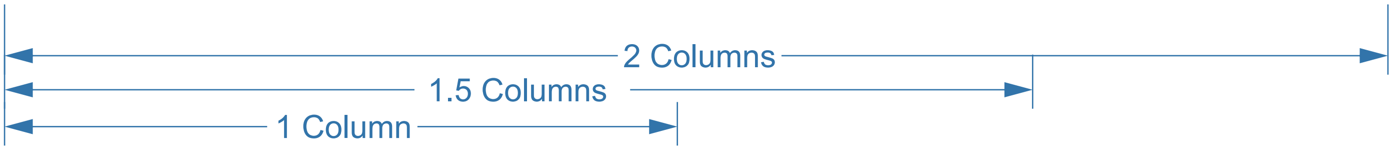


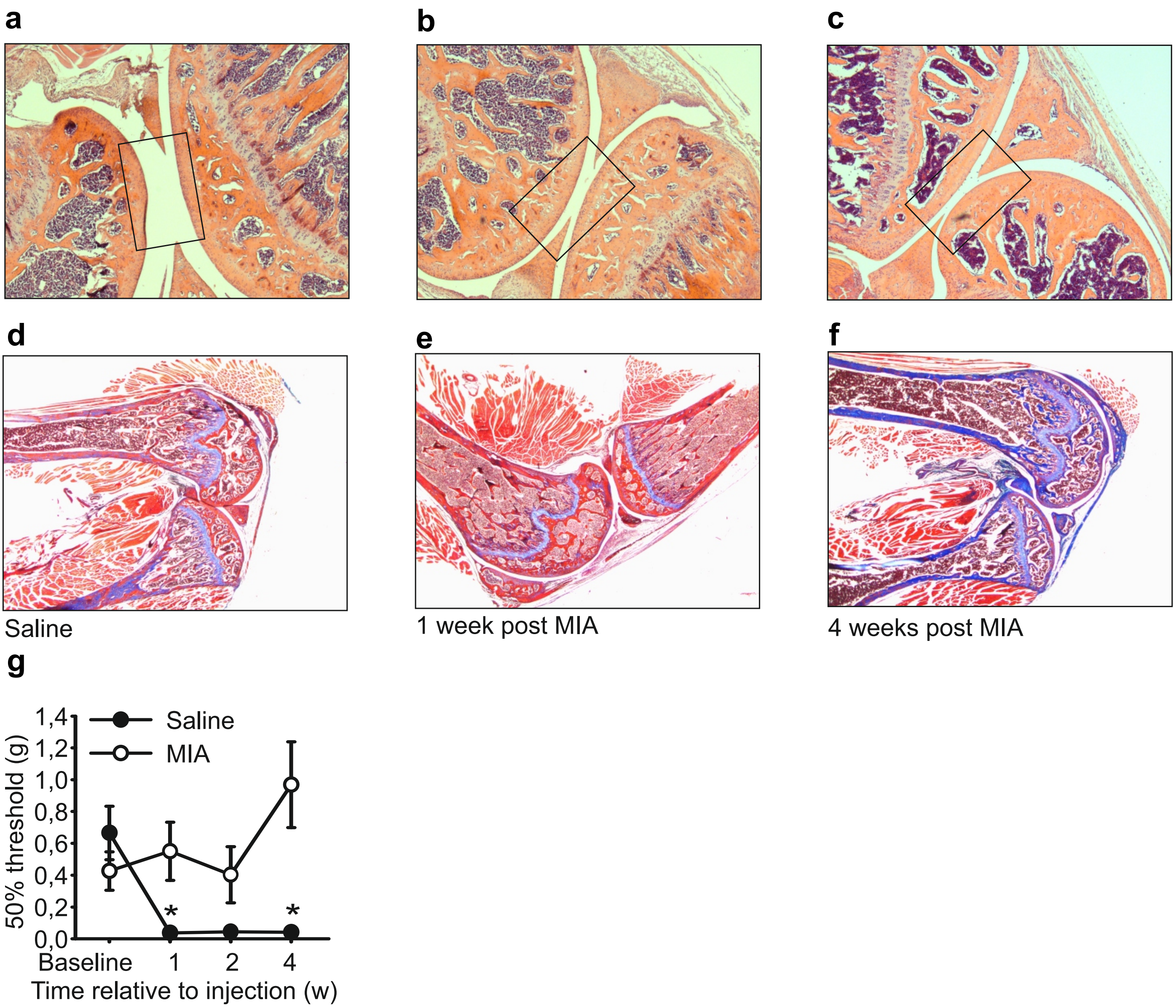


#### Supplementary Figure 1

### Suppl fig 2

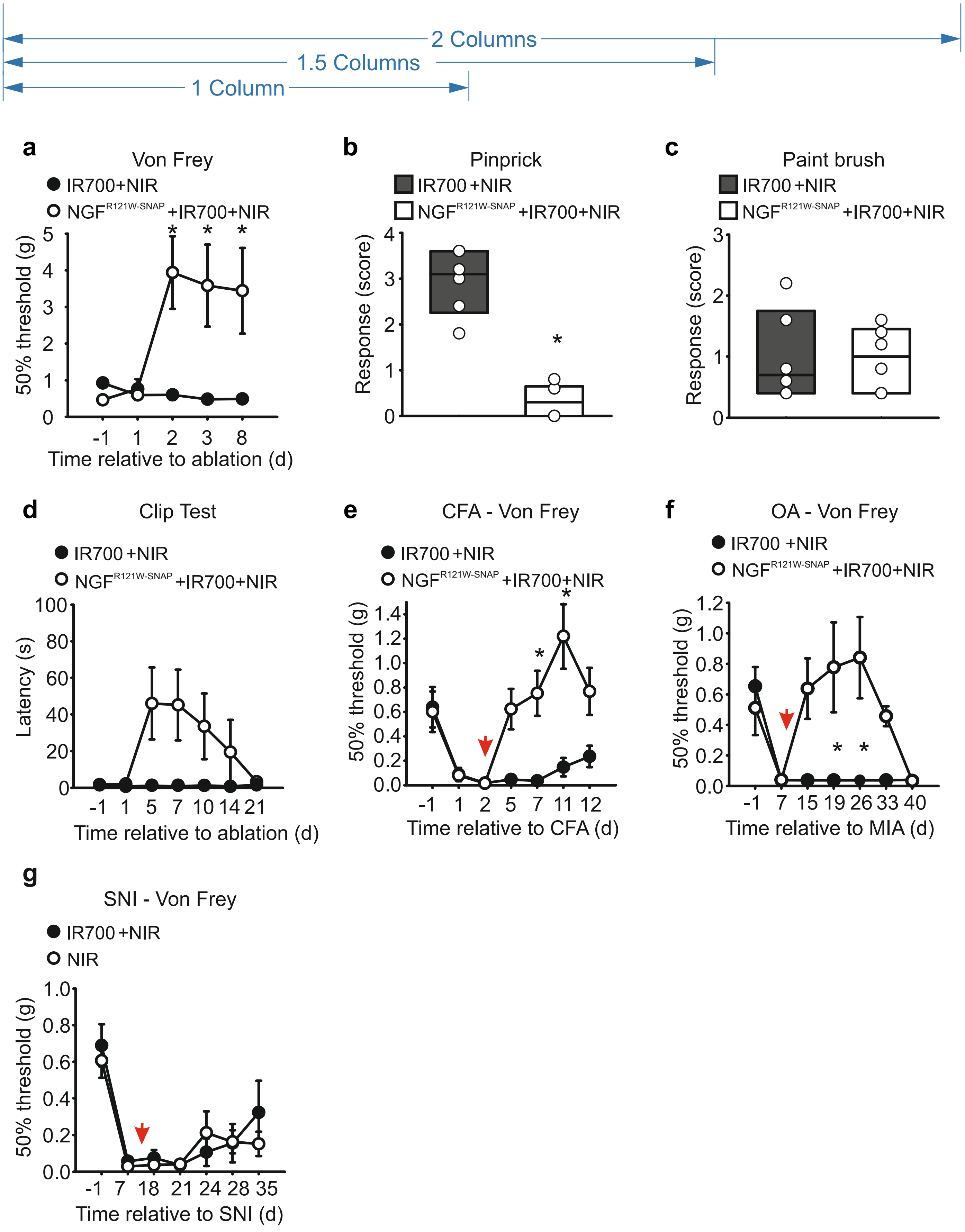


#### Supplementary Figure 2
